# Network-based meta-analysis maps stage-dependent molecular programs in MASLD through MASLD-META NETWORK application

**DOI:** 10.64898/2026.08.26.747338

**Authors:** Eren Kumak, Thomas Darde, Ozlen Konu

## Abstract

Metabolic dysfunction-associated steatotic liver disease (MASLD), the leading cause of chronic liver pathologies worldwide, represents a growing clinical burden. Its diagnosis remains reliant on liver biopsy that limits early detection and the ability to capture molecular changes across disease progression. A systematic understanding of stage-dependent gene expression changes is essential to identify biomarkers and effectively characterize disease mechanisms. Therefore recent studies provided databases for searching genes as well as prediction of multi-gene signatures for disease progression. However, there is still a need for interactive and comprehensive meta-analysis of datasets of MASLD patients with available histological metadata. Herein, we performed a meta-analysis of RNA-seq datasets using NAFLD Activity Score (NAS; n = 897) and fibrosis stage (n = 856) upon conducting pairwise comparisons across histological stages and identified differentially expressed genes associated with disease progression. Most importantly, we provide our findings via a dedicated web server, the MASLD-META NETWORK (https://masld.scilicium.com), enabling users to interactively explore meta-analysis results across diverse network modalities. In addition, we characterized gene expression dynamics across increasing disease stages to identify consistent progression-associated pathways using Louvain clustering. Network-based parameters such as centrality in combination with meta-analysis scores further highlighted central genes and pathways implicated in disease mechanisms. Accordingly, MASLD-META NETWORK enabled an integrative reassessment of recently published gene signatures, identifying COL1A1, COL3A1, THBS2, FBLN5, and PDGFA as the most central genes, and SULF2, MMP14, IL32, GPNMB, and COL3A1 as candidate markers of earlier transcriptional alterations. Network analysis of MASLD associated biological modules further identified LAMA2 and LAMA3 as previously unrecognized central candidate targets.

## Introduction

Metabolic dysfunction-associated steatotic liver disease (MASLD) is an increasingly prevalent disorder that can lead to serious hepatic complications, including cirrhosis and hepatocellular carcinoma (HCC) and systemically cardiovascular disease (CVD) and chronic kidney disease (CKD). MASLD imposes a substantial health and economic burden, particularly alongside obesity and type 2 diabetes (T2D), with an estimated global prevalence of 38% among adults projected to rise to 55.4% by 2040 (Miao et al., 2024). The progression of the disease involves multiple stages, extending from simple steatosis through metabolic dysfunction-associated steatohepatitis (MASH), cirrhosis, and HCC, with pathogenic events occurring in parallel in a multifactorial fashion. This progression reflects the combined effects of metabolic disturbances, hepatic lipid accumulation, inflammatory responses, and activation of fibrogenic pathways within the liver microenvironment (Li et al., 2024).

Currently, MASLD diagnosis involves abdominal ultrasonography, while liver biopsy remains the gold standard for diagnosing MASH and staging fibrosis. Clinical staging of the disease incorporates (a) a five-tier fibrosis score that ranges from F0 (absence of fibrosis) to F4 (cirrhosis), and (b) a composite index from 0 to 8, referred to as the NAFLD Activity Score (NAS), which evaluates steatosis, hepatocyte ballooning, and lobular inflammation (Kleiner et al., 2005). Treatment strategies primarily focus on behavioral modifications, including dietary changes and physical activity while only two pharmacotherapies have received approval from the US Food and Drug Administration (FDA), namely resmetirom and semaglutide (Tilg et al., 2026). However, these drugs target only noncirrhotic MASH and stage F2–F3 fibrosis. Addressing patients in both early and advanced stages of the disease remains challenging, not only from a therapeutic perspective but also in terms of diagnosis. Previous literature highlights the underdiagnosis of patients at early and late stages, driven by limited screening strategies and difficulties in identifying early symptoms (Eskridge et al., 2023). The invasive and costly nature of liver biopsy also contributes to this challenge, as it carries a small but clinically relevant risk of complications such as acute bleeding (Tilg et al., 2026). Although several machine learning based gene signatures have been proposed for disease stage prediction, each was developed and evaluated within a limited set of cohorts, and no study has systematically compared the available models and gene panels across all publicly available datasets with histological scores (Govaere et al. 2020; Kamzolas et al., 2026).

Meta-analysis is a powerful tool for biomarker discovery that allows multiple studies to be analyzed together, increasing the power to detect differentially expressed genes in transcriptomic datasets (Rau et al., 2014). This approach is especially useful when studies include comparable metadata, such as histological grading, which can be linked to disease progression. There have been several previous attempts to perform transcriptomic meta-analyses of MASLD. Govaere et al. (2020) as one of the earliest examples, analyzed 403 samples with respect to both the fibrosis stage and NAFLD Activity Score (NAS). This work was followed by Kendall et al. (2023), who analyzed a larger cohort of 707 samples and incorporated histological scores together with demographic metadata such as age and gender. Kendall et al. (2023) also developed an interactive web server, SteatoSITE, allowing researchers to explore the meta-analysis results through multiple visualization approaches. More recently, a LiRNA webserver was developed that enables gene-gene meta-correlation queries across more comprehensive liver datasets (Alvarez Sola et al., 2026).

Other studies have conducted meta-analyses using datasets that lacked detailed histological metadata. For example, Wang et al. (2021) and Wen et al. (2021) analyzed single-cell transcriptomic datasets which, despite lacking detailed histological staging information, provided higher cellular resolution compared with bulk transcriptomic analyses. Additional meta-analysis studies include those by Piras and DiStefano (2024), Rusu et al. (2025), and Cheng et al. (2024), that analyzed cohorts of 1,058, 458, and 1,304 samples, respectively, although these studies also lacked detailed histological annotations. Cheng et al. (2024) further differed from other analyses by introducing a pseudotime framework modeling disease progression from normal liver to MASLD and MASH. In addition, similar to the SteatoSITE platform developed by Kendall et al., they developed an interactive web server, MegaMASLD, for exploring the results of their meta-analysis (Cheng et al., 2024).

Although some of these above-mentioned studies compared expression differences across fibrosis stages, many did not apply analytical frameworks specifically designed to identify consistent expression trends along ordered disease progression. Earlier transcriptomic studies commonly relied on broad diagnostic categories or comparisons between high and low NAS groups, limiting the resolution of molecular changes across histological severity. Kamzolas et al. (2026) recently addressed this limitation by reconstructing a continuous molecular trajectory; and applying sliding window analysis, WGCNA, regulatory network construction, and network propagation characterized transcription factors and pathways associated with MASLD progression. However, their reference trajectory and regulatory network were derived from two integrated discovery cohorts rather than from a formal meta analysis of all eligible datasets with histological annotations. In addition, although their framework prioritized network associated genes and pathways, it did not provide an interactive platform for identifying genes with the highest centrality within individual biological modules. Moreover, no tool exists that can integrate user supplied gene sets from independent human and cross species studies for comparison with meta-analysis results. Thus, a comprehensive meta analytic framework that combines broad histological cohort coverage, stage resolved pathway module analysis, module specific gene centrality, and interactive external data integration remains lacking.

Here, we present a meta-analysis of nine publicly available transcriptomic datasets comprising liver biopsy samples from 897 MASLD patients. By integrating datasets with available fibrosis stage and NAS annotations, we systematically characterized and statistically annotated gene expression changes across disease progression. We identified genes showing consistent expression trends across disease stages and evaluated their concordance across multiple analytical approaches. We further examined the functional context of the highest-ranking genes through pathway enrichment and network-based analyses. Our analyses identified central genes associated with MASLD progression, including those within individual biological modules, and characterized the specific stages at which these modules emerged across increasing histological severity. To facilitate exploration of these results, we developed an interactive web platform called MASLD-META NETWORK that enables visualization of pathway–pathway and pathway–gene networks, including stage-dependent and sex-specific network views. The platform also allows users to upload independent gene sets, integrate expression direction, calculate module specific centrality metrics, and map mouse and zebrafish gene lists to their human orthologs before making comparisons. Moreover, the user upload and network pruning functionalities of MASLD-META NETWORK were used to evaluate previously proposed predictors of disease progression. Among the existing predictors, COL1A1, COL3A1, THBS2, FBLN5, and PDGFA were prioritized as the most central genes, whereas SULF2, MMP14, IL32, GPNMB, and COL3A1 emerged as candidate markers of earlier transcriptional alterations.

## Method

### Dataset acquisition

Liver biopsy datasets used in this meta-analysis, each annotated with a NAFLD Activity Score, were retrieved from the Gene Expression Omnibus (GEO) database. The raw count data for each study, except GSE225740 and GSE207310, were obtained by downloading the “Series RNA-seq raw counts matrix” provided by the NCBI-generated data option of GEO. For datasets lacking NCBI-generated data, we utilized the author-provided raw counts directly available within the GEO accession records. The metadata of each study was obtained using the getGEO function provided in the GEOquery package in R. (Davis & Meltzer, 2007). These datasets included GSE135251, GSE193066, GSE225740, GSE130970, GSE192959, GSE207310, GSE162694, GSE174478, and GSE185051.

**Table 1.** *Distribution of NAFLD Activity Score (NAS) Across Datasets.* The table presents the number of patients corresponding to each NAS value. The first column is designated for accession numbers of datasets, while the final column indicates the total number of samples per study. The bottom row summarizes the total number of patients per NAS value across all studies.

|  | NAFLD Activity Score (NAS) |  |  |  |  |  |  |  |  |  |
| --- | --- | --- | --- | --- | --- | --- | --- | --- | --- | --- |
| Accession | 0 | 1 | 2 | 3 | 4 | 5 | 6 | 7 | 8 | Total |
| GSE135251 | 10 | 11 | 21 | 26 | 38 | 47 | 37 | 18 | 8 | 216 |
| GSE193066 | 0 | 4 | 10 | 71 | 34 | 20 | 12 | 11 | 2 | 164 |
| GSE225740 | 5 | 6 | 15 | 14 | 18 | 18 | 12 | 5 | 0 | 93 |
| GSE130970 | 4 | 5 | 9 | 18 | 16 | 18 | 8 | 0 | 0 | 78 |
| GSE192959 | 1 | 4 | 10 | 12 | 9 | 11 | 1 | 0 | 0 | 48 |
| GSE207310 | 4 | 1 | 7 | 5 | 6 | 5 | 0 | 1 | 1 | 30 |
| GSE162694 | 32 | 12 | 9 | 11 | 13 | 19 | 12 | 9 | 0 | 117 |
| GSE174478 | 0 | 1 | 11 | 9 | 17 | 36 | 17 | 3 | 0 | 94 |
| GSE185051 | 7 | 2 | 1 | 10 | 19 | 12 | 5 | 1 | 0 | 57 |
| Total | 63 | 46 | 93 | 176 | 170 | 186 | 104 | 48 | 11 | 897 |

**Table 2.** *Metadata Summary of Each Dataset.* The first column lists the GEO accession numbers for each dataset. Additional columns provide key metadata, including sample size, age distribution, sex distribution, and fibrosis scores where available.

| Accession | Age Range | Mean Age ± SD | Sex (M/F) | Fibrosis Score (F0–F4) |
| --- | --- | --- | --- | --- |
| GSE135251 | - | - | - | 46 / 48 / 54 / 54 / 14 |
| GSE193066 | (20 - 83) | 53.2622 ± 13.2215 | 95 / 69 | 6 / 45 / 71 / 41 / 1 |
| GSE225740 | - | - | - | 27 / 26 / 15 / 13 / 6 / 6 (NA) |
| GSE130970 | (19 - 80) | 50.5641 ± 12.20559 | 30 / 48 | 25 / 28 / 9 / 14 / 2 |
| GSE192959 | (56 - 84) | 71.25 ± 5.91608 | 16 / 32 | 1 / 1 / 3 / 16 / 27 |
| GSE207310 | - | - | 3 / 27 | - |
| GSE162694 | (18 - 72) | 44.98291 ± 13.04765 | 31 / 86 | 35 / 30 / 27 / 8 / 12 |
| GSE174478 | (19 - 92) | 55.14894 ± 15.73774 | 49 / 45 | 7 / 29 / 23 / 24 / 11 |
| GSE185051 | (9 - 68) | 16.57895 ± 10.68535 | 32 / 25 | 11 / 37 / 4 / 5 / 0 |

### NAS Grouping

PERMANOVA was used to group samples according to the NAS, to maximize inter-group variability and minimize intra-group variability while ensuring balanced group sizes within each dataset. For each study, PERMANOVA, implemented using the vegan R package, was first applied to the full dataset using the NAS and variance-stabilizing transformed (VST) counts, obtained via the DESeq2 package, to assess overall group differences (Love, Huber, & Anders, 2014; Oksanen et al., 2022). Then, pairwise PERMANOVA comparisons were performed between each NAS value to evaluate specific group-level distinctions. The resulting p-values were corrected for multiple testing using the Benjamini-Hochberg method (Benjamini & Hochberg, 1995). Grouping decisions were made using majority voting across datasets. Specifically, for datasets in which the initial PERMANOVA (applied to the full dataset) indicated a significant overall difference (adjusted p < 0.05), pairwise comparisons were assessed. The proportion of significant adjusted p-values (< 0.05) for each pairwise comparison was calculated across studies. NAS scores were then grouped based on pairs showing a lower frequency of significant differences in this majority vote, while also ensuring comparable group sizes across datasets (Supplementary Figure 1).

### Regression Analysis

A linear model was fit to NAS groupings, fibrosis scores and VST counts in each study using the default lm() function of R. Then, for each study, adjusted p-values were calculated using the Benjamini-Hochberg method. Genes with adjusted p-values below 0.05 were identified in each dataset. For these, average R^2^, mean slope, total frequency across datasets, and the number of datasets in which the slope was positive or negative were recorded.

### Gene Disease Patterns

For each individual study, the gene disease patterns were identified for the NAS grouping and fibrosis scores using the workflow presented in Aibar et al. (2016). First, GAMMA rank correlations were calculated to get genes with expression profiles correlating with the progression of the NAS values. Next, a Self Organizing Map (SOM) was applied to cluster genes with specific expression patterns characterized with increases in NAS values. After creating code and counts plots for each study, the number of occurrences of each gene across studies, directionality of these occurrences, and associated expression patterns within individual studies were also recorded.

### Differential Gene Expression Analysis

#### Individual DGEA

In each dataset, DESeq2 was performed individually according to NAS groupings and fibrosis scores. For filtering of counts, genes that are expressed (≥ 10 reads) in at least as many samples as the smallest group were kept. Then, pair-wise comparisons of each NAS group as well as fibrosis score were performed.

Meta-p-values were calculated using the metaRNASeq library (Rau et al., 2014). As the method assumes that p-values under the null hypothesis follow a uniform distribution, histograms of p-values from each study were visually examined. Additionally, the Wasserstein distance was calculated between the p value distributions corresponding to the groups with the highest and lowest NAS values or fibrosis scores, depending on the analysis. The distance observed between these most extreme groups was used as the inclusion threshold, and only studies with a Wasserstein distance less than or equal to this value were retained for meta-analysis, thereby selecting studies with p value distributions more consistent with the null hypothesis.

After meta-p-values were obtained using both the inverse normal and Fisher methods, filtering was performed to keep genes with a p-value below 0.05 in both of the meta-p-values and with consistent logFC directions across all individual analyses. Following filtering, the logFC values from each study, as well as the median and average logFC, were recorded for each gene.

#### Global DGEA

After merging the metadata and raw counts for each study, batch correction was performed using ComBat-seq, with ‘study’ specified as the batch variable and NAS grouping or fibrosis score used as the biological condition (Zhang, Parmigiani, & Johnson, 2020). Following batch adjustment, differential gene expression analysis was conducted on the prefiltered counts, retaining only genes with counts ≥ 10 in at least 95% of the samples. After getting the pair-wise comparisons of each NAS group or fibrosis score, only genes with adjusted p-value below 0.05 were kept.

### Scoring

In order to ensure the consistency of each gene across different types of analyses, a frequency-based scoring system was applied. In this system, the total score for each gene was defined as the sum of its occurrences across individual studies in the regression analysis and gene–disease pattern workflow, as well as its inclusion in individual and global differential gene expression comparisons. Finally, 5% and 1% cutoffs were applied to the total score, resulting in the selection of 951 and 163 genes, as significant top candidates, respectively for NAS groupings, and selection of 1028 and 240 genes, respectively for fibrosis scores.

### Over-Representation Analysis

Over-representation analyses were performed using the enricher() function from the clusterProfiler package (Wu et al., 2021). P-values were adjusted using the Benjamini-Hochberg method, and only results with a q-value below 0.05 were retained. Gene sets were obtained via the msigdbr() function from the msigdbr package (Dolgalev, 2022). Transcription factor–gene sets were sourced from the DoRothEA, filtered for confidence levels ‘A’, ‘B’, and ‘C’ using the dorothea package during over-representation analysis (Garcia-Alonso et al., 2019). Visualization of over-representation results was done using functions from the enrichplot package (Yu, 2023).

### Genotype-Tissue Expression Project

To evaluate the cross-tissue expression profiles of NAS- and fibrosis-associated candidate genes, bulk RNA-sequencing data were obtained from the Genotype-Tissue Expression (GTEx) project (v10 release; gene-level median transcripts per million [TPM]). Raw expression matrices were preprocessed to truncate Ensembl identifier version suffixes, and TPM values for redundant Ensembl IDs were averaged. Identifiers were strictly mapped to official HGNC gene symbols using the org.Hs.eg.db R package; unmapped genes and ambiguous many-to-one alignments were excluded to prevent cross-annotation artifacts.

For each candidate gene, median TPM values were queried across all GTEx tissues to establish a multi-tissue expression profile. Genes were subsequently stratified by their relative hepatic expression into one of four distinct categories: (1) highest expression in the liver (maximum cross-tissue TPM localized to the liver, encompassing liver-exclusive transcripts); (2) expressed in the liver (hepatic TPM > 0, but maximal TPM observed in an extrahepatic tissue); (3) not expressed in the liver (hepatic TPM = 0); or (4) undetected within the filtered GTEx dataset. All transcriptomic data parsing, ID mapping, and categorical assignments were executed in R.

### MASLD-META NETWORK Application

An interactive web application for visualization and integrative analysis of meta-analysis results was developed using R Shiny (Chang et al., 2026). Network construction, visualization, and graph-based metric calculations were implemented using the igraph and visNetwork packages (Csárdi et al., 2026; Almende B.V. et al., 2022).

Protein–protein interaction networks were obtained from STRING v12.0 (Szklarczyk et al., 2025), retaining interactions with confidence scores ≥ 0.4 (medium confidence). Gene-level meta-analysis scores were integrated with network topology by combining standardized meta-analysis scores with network centrality measures. Combined gene scores were calculated using user-adjustable weighting of meta-analysis–derived scores and selected centrality metrics.

Over-representation analysis results derived from MSigDB gene set collections and DoRothEA transcription factor target sets were integrated into pathway–pathway gene overlap networks, in which edges represent similarity between gene sets based on shared gene membership (Dolgalev, 2022; Garcia-Alonso et al., 2019). Network pruning can be performed using either Jaccard similarity coefficients (Keskus et al., 2026) or the absolute number of shared genes between pathways. Thresholds for network pruning and statistical significance filtering (q-value cutoffs) are user-adjustable to allow flexible control of network density and interpretability.

Pathway similarity networks can be partitioned into modules representing groups of functionally related pathways using multiple community detection algorithms, including Louvain, edge betweenness, and label propagation clustering (Csárdi et al., 2026). In MASLD-META NETWORK application gene–pathway bipartite networks were constructed to identify genes shared across multiple related pathways, enabling identification of central genes within pathway modules.

Network centrality can be calculated using multiple graph-based metrics, including degree centrality, closeness centrality, betweenness centrality, and PageRank, allowing users to evaluate gene importance under different topological assumptions (Csárdi et al., 2026). Combined gene scores were calculated using adjustable weighting between meta-analysis scores and network-derived centrality measures, enabling flexible prioritization strategies.

The platform supports gene-level filtering of networks based on differential expression direction. Users may upload custom gene lists together with log fold-change values to enable filtering for concordantly regulated genes across their study and meta-analysis. Cross-species compatibility is supported through integration with DIOPT, enabling mapping of homologous genes from *Mus musculus* and *Danio rerio* to human orthologs for downstream network analysis (Hu et al., 2011; Hu et al., 2026). To maximize mapping accuracy, we retained only the orthologs with the best DIOPT scores for our final results in both directions.

The application additionally provides stage-dependent network visualizations, enabling comparison of pathway organization across increasing histological severity for both NAS and fibrosis stage analyses. Sex-stratified network representations are also available, allowing comparison of pathway enrichment patterns derived from meta-analyses performed separately for each sex.

A transcriptome browser module is also included to enable visualization of gene-level meta-analysis results across disease stages for all genes and not just top scoring ones. Users may query individual genes of interest or upload gene lists to visualize stage-dependent changes in NAS and fibrosis analyses.

The MASLD-META web application is publicly available at https://masld.scilicium.com. The source code used to generate the application interface will be made publicly available upon publication.

## Results

### Identification of Top-Scoring Genes Associated with MASLD Progression

After filtering genes based on the 0.95 quantile of the total score, 951 genes were selected from 14,713 for NAS-related analysis, while analysis with respect to fibrosis score yielded 1,028 genes from 20,119 (Supplementary Data 1). Among the 951 NAS-associated genes, 390 had not been previously reported in PubMed when queried using terms including “NAFLD”, “NASH”, “MASLD”, “MASH”, “nonalcoholic fatty liver disease”, “metabolic dysfunction-associated steatotic liver disease”, “steatohepatitis”, “fatty liver”, “hepatic steatosis”, “liver steatosis”, “liver fibrosis”, and “hepatic inflammation” using the easyPubMed package (Fantini, 2019). A similar analysis identified 421 such genes among the 1,028 fibrosis-associated genes. According to the Genotype-Tissue Expression (GTEx) Project, 91 NAS-associated top-scoring genes showed highest expression in the liver, compared with 84 genes for fibrosis-associated top-scoring genes (Lonsdale et al., 2013). The majority of genes in both groups (853 and 924, respectively) were also expressed in the liver, although not at the highest level. The remaining genes were either not expressed in the liver or not detected in GTEx.

### Network Topology Identifies Core Interaction Hubs Across MASLD Progression

To identify genes occupying central positions within disease-relevant interaction networks, meta-analysis–derived gene scores were integrated with protein–protein interaction topology using the MASLD-META NETWORK application. Combined centrality scores were calculated by weighting meta-analysis scores and STRING-derived degree centrality equally, enabling systematic identification of shared and stage-specific interaction hubs across NAS and fibrosis (Supplementary Data 2).

#### NAS-associated interaction hubs highlight extracellular matrix remodeling and inflammatory myeloid activation

Genes with the highest combined scores with respect to NAS were predominantly associated with extracellular matrix organization and inflammatory myeloid activation, together with a smaller subset linked to cellular proliferation and hepatocyte identity. The 20 most central genes were *COL1A1*, *COL1A2*, *COL3A1*, *ITGAX*, *ANXA2*, *COL5A1*, *LGALS3*, *THBS2*, *TGFB1*, *SRC*, *TIMP1*, *IL32*, *GPNMB*, *TOP2A*, *SPP1*, *ITGAM*, *ITGB2*, *APOF*, *COL4A2*, and *SQSTM1*. Among these central genes, all except APOF were upregulated. Subnetwork visualization (Fig. 1A) restricted to this subset revealed a densely connected interaction structure, with APOF and IL32 not directly connected to the main component. All 20 central genes had previously been reported in MASLD according to PubMed queries performed using the easyPubMed package.

**Figure 1.**
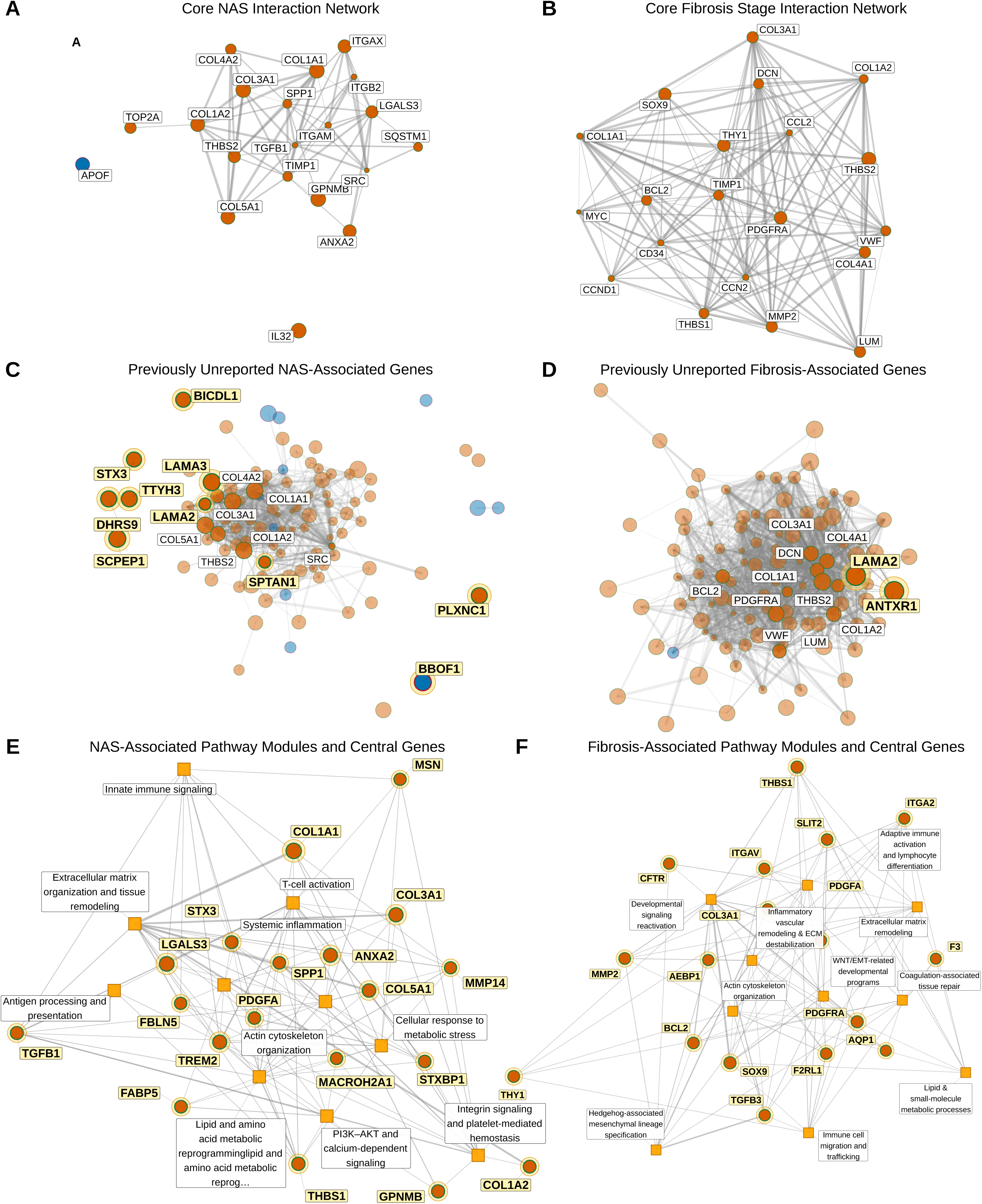
*Network topology identifies central genes and biological modules associated with NAS and fibrosis severity.* **A, B,** STRING protein protein interaction networks of the genes with the highest combined scores in the NAS **(A)** and fibrosis **(B)** analyses. Combined scores integrated standardized meta analysis scores and degree centrality with equal weighting. Node size represents the combined score. Orange and blue nodes indicate upregulated and downregulated genes, respectively. Grey edges represent STRING interactions retained at a confidence score of at least 0.4. **C, D,** Protein protein interaction networks of the 100 highest ranked genes associated with NAS **(C)** and fibrosis **(D)**. Genes not previously associated with MASLD in PubMed searches are outlined, labelled, and highlighted, together with connected genes that are ranked among those with the highest combined scores. Node colour indicates the direction of expression change, as in A and B. **E, F,** Bipartite networks connecting major pathway modules with the most central genes identified in the NAS **(E)** and fibrosis **(F)** analyses. Squares represent biological pathway modules, circles represent genes, and edges indicate the inclusion of a gene within a module. Pathway modules were derived from significantly enriched ontology gene sets with q < 0.05, connected using a Jaccard similarity threshold of 0.2, and grouped using Louvain community detection.

Extension of the analysis to the top 100 ranked genes identified 10 genes not previously associated with MASLD in PubMed: *LAMA3*, *BBOF1*, *SCPEP1*, *DHRS9*, *PLXNC1*, *TTYH3*, *BICDL1*, *LAMA2*, *STX3*, and *SPTAN1*. Among these, *BBOF1* was the only downregulated gene. *LAMA3*, *LAMA2*, and *SPTAN1* were connected to the main high-centrality component, whereas the remaining genes primarily interacted with lower-ranked nodes. Notably, DHRS9 and TTYH3 were connected to each other but did not directly connect to the principal hub structure (Fig. 1C).

#### Fibrosis-associated hubs emphasize mesenchymal and stromal remodeling programs

Central genes identified for fibrosis were strongly enriched for extracellular matrix organization and mesenchymal or stromal cell biology. The 20 highest-ranking genes were *BCL2*, *MMP2*, *COL1A2*, *COL1A1*, *MYC*, *COL3A1*, *LUM*, *THBS2*, *THBS1*, *PDGFRA*, *DCN*, *CCL2*, *TIMP1*, *SOX9*, *CD34*, *THY1*, *VWF*, *COL4A1*, *CCN2*, and *CCND1*. All 20 fibrosis-associated hub genes were upregulated, and subnetwork visualization demonstrated a connected interaction structure (Fig. 1B). Each of these genes had previously been reported in MASLD according to the PubMed queries performed using the easyPubMed package. Analysis of the top 100 ranked genes identified only two genes not previously associated with MASLD in PubMed: *LAMA2* and *ANTXR1*. Both genes were upregulated and connected to the central interaction component (Fig. 1D)

### Pathway Network Architecture Reveals Modular Organization of MASLD Biology

Over-representation analysis of top-scoring genes revealed extensive pathway enrichment for both top-scoring NAS and fibrosis associated genes. Enriched ontology gene sets were integrated using the MASLD-META NETWORK application to identify higher-order functional structure based on gene overlap between pathways. Pathway relationships were defined using a Jaccard similarity threshold of 0.2 to retain robust functional connections, and network structure was partitioned using Louvain community detection to identify groups of biologically related pathways. This analysis identified 1,029 significantly enriched pathways for NAS and 881 for fibrosis (q < 0.05), which were organized into 34 pathway modules in each analysis. Module sizes ranged from 2 to 146 pathways in NAS and from 2 to 120 pathways in fibrosis, indicating comparable modular complexity across disease stages (Supplementary Data 3-4).

#### NAS-associated modules reflect inflammatory remodeling and metabolic stress adaptation

The largest NAS-associated pathway modules corresponded to biological processes related to systemic inflammation, extracellular matrix organization and tissue remodeling, T-cell activation, integrin signaling and platelet-mediated hemostasis, lipid and amino acid metabolic reprogramming, actin cytoskeleton organization, cellular response to metabolic stress, innate immune signaling, PI3K–AKT and calcium-dependent signaling pathways, and antigen processing and presentation (Fig. 1E).

Integration of meta-analysis–derived scores with module-specific network topology identified *COL1A1*, *LGALS3*, *COL3A1*, *ANXA2*, *TREM2*, *COL5A1*, *GPNMB*, *THBS1*, *TGFB1*, *STX3*, *COL1A2*, *FABP5*, *PDGFA*, *STXBP1*, *FBLN5*, *AJUBA*, *MACROH2A1*, *MSN*, *SPP1*, and *MMP14* as the most central genes across NAS-associated modules, all of which were upregulated. Furthermore, MASLD-META NETWORK application allowed calculating centrality metrics for each module separately identifying central genes of each NAS-associated module separately (Supplementary Data 5).

Inspection of pathway enrichment according to the expression direction of their contributing genes revealed distinct regulatory patterns within specific modules (Supplementary Data 5). The lipid and amino acid metabolic reprogramming module included pathways enriched among downregulated genes, particularly those involved in small molecule metabolism and amino acid catabolism, as well as pathways enriched among upregulated genes, including lipid metabolism and lipid derived inflammatory signaling pathways associated with omega 6 polyunsaturated fatty acids. Similarly, within the cellular response to metabolic stress module, pathways related to the positive regulation of carbohydrate metabolism were predominantly enriched among downregulated genes, whereas oxidative stress response and autophagy pathways were enriched among upregulated genes. These findings indicate that distinct stress adaptive and metabolic programs were associated with opposing gene expression patterns across NAS progression. In the remaining modules, pathway enrichment was predominantly driven by upregulated genes.

#### Fibrosis-associated pathway modules indicate reactivation of developmental and mesenchymal programs

Fibrosis-associated pathway modules highlighted biological processes related to reactivation of developmental signaling programs, Hedgehog-associated mesenchymal lineage specification, inflammatory vascular remodeling and extracellular matrix destabilization, adaptive immune activation and lymphocyte differentiation, WNT signaling and epithelial–mesenchymal transition–related developmental programs, lipid and small-molecule metabolic processes, extracellular matrix remodeling, immune cell migration and trafficking, actin cytoskeleton organization, and coagulation-associated tissue repair (Fig. 1F).

Integration of meta-analysis scores with module-specific network topology identified *THBS1*, *SOX9*, *MMP2*, *PDGFRA*, *THY1*, *SLIT2*, *BCL2*, *TGFB3*, *CFTR*, *COL3A1*, *F2RL1*, *PDGFA*, *AEBP1*, *AQP1*, *ITGAV*, *VEGFC*, *FLNA*, *ITGA2*, *ITGA3*, *F3* as the most central genes across fibrosis-associated modules, all of which were upregulated. Similar to NAS-associated modules, the central genes of each individual fibrosis-associated module were also investigated via MASLD-META NETWORK application (Supplementary Data 6). Across fibrosis-associated modules, enriched pathways were predominantly associated with upregulated genes. An exception was observed for the lipid and small-molecule metabolic processes module, in which pathways related to small-molecule catabolic processes and alcohol and hydroxy-compound metabolism were primarily associated with downregulated genes. In contrast, other pathways within the same module, including fatty acid synthesis, glycosaminoglycan synthesis, Golgi apparatus organization, epithelial polarity, transmembrane transport, and nucleotide- and cAMP-related signaling pathways, were enriched among upregulated genes.

### Transcriptional Regulators Underlying MASLD Progression

DoRothEA based over representation analysis identified transcription factors whose targets were enriched among top scoring genes associated with NAS and fibrosis (Supplementary Data 7). *RELA*, *SMAD3*, *RUNX2*, and *TP53* were identified across both histological scores, suggesting that genes associated with MASLD severity were regulated by shared inflammatory, stress responsive, and extracellular matrix related transcriptional programs.

Among NAS associated genes, *RFX5*, *SMAD3*, *SPI1*, *RUNX2*, and *RELA* were significantly enriched. Analysis according to gene expression direction additionally identified *TP53*, *GLI2*, *POU2F2*, and *IKZF1* among upregulated genes, whereas HNF4A was enriched among downregulated genes. Fibrosis associated genes showed broader transcription factor enrichment, including regulators of inflammatory signaling, developmental processes, and cell identity. *ZEB1* was additionally identified among upregulated genes, while *HNF4A* and *CEBPA* were enriched among downregulated genes.

Overall, these results identify candidate transcriptional regulators of genes associated with MASLD severity. The enrichment of inflammatory and developmental regulators among upregulated genes, together with *HNF4A* and *CEBPA* enrichment among downregulated genes, suggests that MASLD progression related gene signatures encompass inflammatory and tissue remodeling programs alongside reduced expression of genes linked to hepatocyte identity.

### Functional Characterization of Genes Associated with NAS and Fibrosis Progression

To evaluate how MASLD progression is reflected across NAS and fibrosis stages, we compared pathway similarity network structures derived from over-representation analyses performed for each histological stage relative to its baseline group. In these networks, edges represent similarity between pathways based on shared genes. For fibrosis, stages F2, F3, and F4 were independently compared against a baseline group consisting of F0 and F1 samples. For NAS, group 0 (NAS 1-2-3) was used as the baseline for comparison with higher NAS groups. Differential gene expression analysis was performed separately within each dataset, followed by meta-analysis to identify consistently dysregulated genes. Pathway relationships were defined using a Jaccard similarity threshold of 0.2, and network structure was partitioned using Louvain community detection to identify groups of biologically related pathways (Supplementary Data 8-9)

#### Stage-specific pathway similarity networks across NAS progression

Comparison of NAS Group 1 (NAS = 3) against the baseline Group 0 (NAS = 0–1–2) identified 46 significantly over-represented pathways (q < 0.05). Network clustering of these pathways resulted in two major modules, both primarily related to immune-associated processes. The first module additionally included over-representation of superoxide metabolic process, whereas the second module was more specifically associated with adaptive immune activation and ERK/MAPK signaling (Fig. 2A). Pathways within both modules were primarily driven by genes that were upregulated in the meta-analysis.

**Figure 2.**
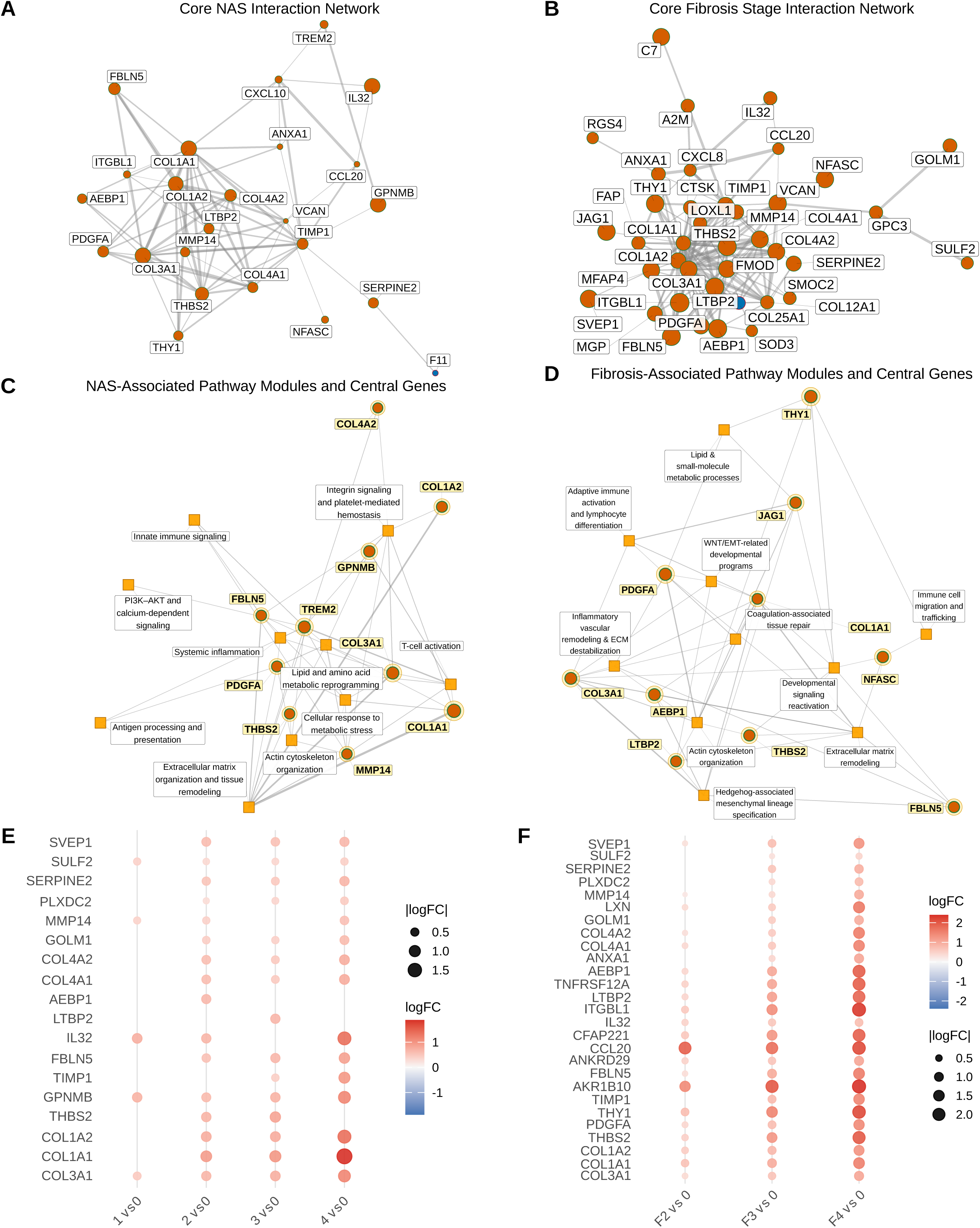
*MASLD-META NETWORK refines previously reported gene panels into stage resolved priorities.* **A, B,** STRING protein protein interaction networks pruned to genes compiled from six previously reported MASLD gene panels and recovered among the top scoring genes in the NAS **(A)** and fibrosis **(B)** analyses. Nodes represent genes, and edges represent STRING interactions retained at a confidence score of at least 0.4. Node size reflects the combined score obtained by equally weighting the standardized meta analysis score and degree centrality. **C, D,** Bipartite networks showing the ten most central published panel genes selected to represent the major biological modules identified in the NAS **(C)** and fibrosis **(D)** analyses. Circles represent genes, squares represent biological pathway modules, and edges indicate the inclusion of a gene within a module. **E, F,** Stage resolved expression profiles of prioritized genes across NAS **(E)** and fibrosis **(F)** comparisons. NAS groups 1, 2, 3, and 4 correspond to NAS 3, 4, 5, and 6 to 8, respectively, and were compared with group 0, comprising NAS 0 to 2. Fibrosis stages F2, F3, and F4 were independently compared with the F0 plus F1 baseline group. Dot size represents the absolute log fold change, and dot colour represents the direction and magnitude of the log fold change. Dots indicate comparisons in which genes were retained as differentially expressed in the meta analysis. logFC values are obtained by getting average logFCs across meta analyzed datasets.

When the disease stage increased to NAS Group 2 (NAS = 4), the number of significantly over-represented pathways increased to 229, which were organized into four modules (Fig. 2B). The two largest modules (n = 56 and n = 52) were primarily associated with extracellular matrix remodeling. Additional modules corresponded to amino acid and nitrogen transfer metabolic processes (n = 30), actin cytoskeleton organization (n = 13), and coagulation-associated tissue repair (n = 18). Pathways within the amino acid metabolism and coagulation-associated modules were primarily driven by downregulated genes, whereas the actin cytoskeleton module was associated with upregulated genes. In contrast to NAS Group 1, immune-related modules were not observed among the modules identified for NAS Group 2.

Increasing the comparison to NAS Group 3 (NAS = 5) resulted in 275 significantly over-represented pathways (q < 0.05) (Fig. 2C). The extracellular matrix module remained present (n = 64). In addition, a module combining ERK/MAPK signaling, platelet biology pathways, and general adhesion-related processes was identified (n = 37), while pathways related to regulation of wound healing and coagulation were no longer observed. Modules corresponding to amino acid and nitrogen transfer metabolic processes also continued to be present (n = 37). Two additional modules were identified that were primarily composed of pathways associated with downregulated genes in the meta-analysis, corresponding to carbohydrate metabolism and insulin-response regulation (n = 22) and lipid, bile acid, and steroid metabolic remodeling (n = 24).

As the disease stage further increased to NAS Group 4 (NAS = 6-7-8), the number of significantly over-represented pathways increased to 830 (q < 0.05) (Fig. 2D). Modules primarily associated with upregulated genes included extracellular matrix remodeling (n = 95), immune cell recruitment, activation, and cytokine signaling, including both myeloid- and T-cell–related processes together with ERK signaling (n = 107), and actin–integrin adhesion and wound-repair remodeling (n = 72). In addition, modules not previously observed at earlier NAS stages emerged, including apical actomyosin and hepatobiliary epithelial polarity (n = 80) and complement and humoral immune activation (n = 79).

In contrast, none of the modules primarily composed of pathways associated with downregulated genes were newly introduced. These included lipid, bile acid, and steroid metabolic remodeling (n = 68), carbohydrate metabolism and insulin-response regulation (n = 67), and amino acid metabolism and nitrogen transfer metabolic processes (n = 53), all of which were also observed at earlier stages. Additional smaller modules were also identified, including a PI3K–AKT and MAPK signaling module (n = 19) and a hepatobiliary lipid and bile acid transport module (n = 18).

#### Stage-specific pathway similarity networks across fibrosis progression

Comparison of F2 against baseline fibrosis (F0+F1) resulted in 467 significantly over-represented pathways (q < 0.05). Network clustering identified 10 primary modules in this comparison (Fig. 2E). Among modules mainly driven by genes upregulated in the meta-analysis were extracellular matrix remodeling (n = 41), cell adhesion and homeostasis (n = 28), and mitotic cell cycle progression (n = 27). In contrast, modules primarily composed of pathways associated with downregulated genes were more abundant and included Ribosomopathy-associated pathways (n = 69), oxidative phosphorylation and respiratory chain (n = 53), amino acid and fatty acid oxidative catabolism (n = 50), hepatic P450-dependent detoxification together with lipid and steroid metabolism (n = 45), lipoprotein remodeling and cholesterol efflux (n = 35), oxidative stress and detoxification (n = 27), and ribosome biogenesis and mitochondrial translation (n = 23).

Further increasing the stage to F3 against baseline increased the number of significantly over-represented pathways to 877 (q < 0.05) (Fig. 2F). At this stage, previously identified metabolism-related modules observed in F2 were largely merged into a broader oxidative intermediary metabolism module that included fatty acid metabolism, amino acid metabolism, and one-carbon metabolism related pathways (n = 58). The previously identified lipoprotein remodeling and cholesterol efflux module decreased in size (n = 23), and modules corresponding to ribosome biogenesis and mitochondrial translation as well as Ribosomopathy-associated pathways were no longer observed. Compared with F2, this comparison contained a greater number of modules composed of pathways associated with genes upregulated in the meta-analysis, including extracellular matrix remodeling (n = 86), adaptive immunity (n = 51), developmental morphogenesis and stem cell differentiation (n = 53), cell adhesion and hemostasis (n = 29), coagulation-associated tissue repair (n = 29), receptor tyrosine kinase signaling (MAPK/ERK and PI3K–AKT) (n = 28), solute and ion transport (n = 26), actin cytoskeleton organization (n = 18), and mitochondrial apoptosis together with p53-mediated DNA damage response (n = 12).

Visualization of the stage-dependent network for F4 compared with baseline resulted in 1022 significantly over-represented pathways (q < 0.05) (Fig. 2G). Similar to the F3 comparison, F4 included a general oxidative intermediary metabolism module (n = 119), which did not introduce additional metabolic categories and consisted of pathways related to fatty acid metabolism and β-oxidation, amino acid metabolism, alcohol and aldehyde metabolism, oxidoreductase activity, lipid-derived signaling molecules and steroid metabolism, and one-carbon and folate metabolism. Lipoprotein remodeling, with a reduced cholesterol-related signature, continued to be present (n = 30). This comparison also included a general systemic metabolic dysfunction phenotype module (n = 53), consisting of pathways associated with abnormal circulating metabolites, including circulating lipids, blood glucose, nitrogen compounds, hepatobiliary dysfunction pathways, and abnormalities in vitamin metabolism.

Modules also continued to include extracellular matrix remodeling (n = 57), developmental morphogenesis and stem cell differentiation (n = 63), actin cytoskeleton organization (n = 41), receptor tyrosine kinase signaling (MAPK/ERK and PI3K–AKT) (n = 31), coagulation-associated tissue repair (n = 26), and cell adhesion and hemostasis (n = 24). In contrast to F3, which included a single adaptive immunity module, this comparison contained multiple immune-related modules, including adaptive immunity (n = 63), an additional immune-related module including myeloid-associated signaling primarily related to macrophage biology (n = 49), and a module related to immune cell migration (n = 21).

### MASLD-META NETWORK refines published gene panels into stage-resolved priorities

Six previous meta analyses had proposed gene panels associated with MASLD or the prediction of disease severity. To demonstrate the utility of the publicly available MASLD-META NETWORK application, we compiled the genes reported across these panels and used the resulting set of 156 genes to prune the NAS and fibrosis associated networks. The application enables users to examine gene level meta analysis results while simultaneously assessing the position and centrality of each gene within protein protein interaction networks and biological pathway modules.

Agreement among the published panels was limited. Only five of the 156 genes were reported by more than one study, and no gene was identified by more than two studies. Of the 156 genes, 76 were recovered in our top scoring gene lists. Among these, 20 ranked highly in both the NAS and fibrosis analyses, whereas 37 were specific to the fibrosis analysis and 11 were specific to the NAS analysis. Eight of the recovered genes also ranked among the most central genes in either the protein protein interaction networks or the biological pathway modules.

Pruning the protein protein interaction networks to the published panel genes revealed a connected cluster in both the NAS and fibrosis networks. When meta analysis scores and degree centrality were weighted equally, the collagen genes COL1A1, COL1A2, COL3A1, COL4A1, and COL4A2, together with THBS2 and TIMP1, ranked among the most prominent genes within both clusters (Fig. 3A and Fig. 3B). We further reduced the published panels to the ten most central genes required to represent all major biological modules identified for each histological score. Five genes, COL1A1, COL3A1, THBS2, FBLN5, and PDGFA, were retained in both the NAS and fibrosis module networks (Fig. 3C and Fig. 3D), indicating that they connect multiple biological processes across complementary measures of MASLD severity.

**Figure 3.**
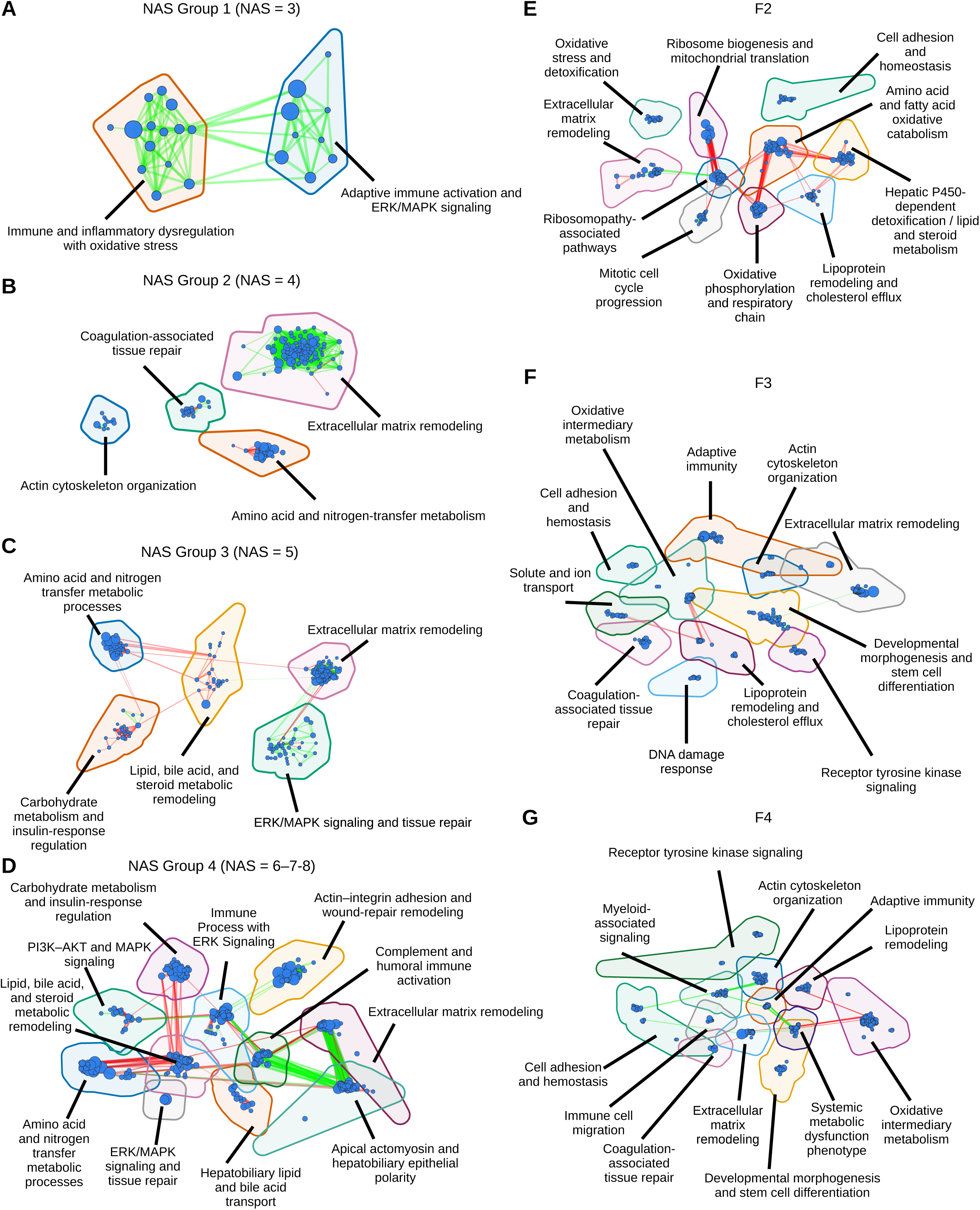
Stage resolved pathway similarity networks reveal the emergence and reorganization of biological modules across MASLD progression. **A to D,** Pathway similarity networks derived from differential expression meta analyses comparing NAS group 1, defined as NAS 3 **(A)**, NAS group 2, defined as NAS 4 **(B),** NAS group 3, defined as NAS 5 **(C),** and NAS group 4, defined as NAS 6 to 8 **(D),** with the baseline group 0, defined as NAS 0 to 2. **E to G**, Pathway similarity networks derived from comparisons of fibrosis stages F2 **(E)**, F3 **(F)**, and F4 **(G)** with the baseline group comprising F0 and F1. Nodes represent significantly over represented pathways with q < 0.05, and node size reflects the number of contributing genes. Edges connect pathways with a Jaccard similarity of at least 0.2 based on shared gene membership. Pathways were partitioned using Louvain community detection, with colored contours delineating biological modules. Module labels summarize the predominant biological processes represented within each community.

The transcriptome browser implemented in MASLD-META NETWORK further enabled us to determine the pairwise stage comparisons in which these prioritized genes were differentially expressed. Among genes that ranked highly for both histological scores or occupied central positions within the networks, most were already differentially expressed in the F2 versus F0 plus F1 comparison (Fig. 3F). By contrast, only SULF2, MMP14, IL32, GPNMB, and COL3A1 were differentially expressed in the earliest NAS comparison, which contrasted NAS group 1, defined as NAS 3, with NAS group 0, defined as NAS 0, 1, or 2 (Fig. 3E). These genes may therefore capture molecular alterations that precede the broader transcriptional changes observed at more advanced disease stages.

Collectively, MASLD-META NETWORK refined previously reported MASLD gene panels by integrating evidence across studies, meta-analysis performance, network centrality, biological module coverage, and the timing of differential expression across disease stages. This approach prioritized genes with complementary relevance to MASLD progression, including COL1A1, COL3A1, THBS2, FBLN5, and PDGFA as central genes shared across the NAS and fibrosis networks, together with SULF2, MMP14, IL32, GPNMB, and COL3A1 as candidates associated with earlier transcriptional alterations. These genes therefore represent the strongest priorities within the original panel of 156 genes for future investigation.

## Discussion

In this study, we integrated transcriptomic data from nine independent liver biopsy cohorts to characterize molecular changes across histologically defined MASLD progression. By combining differential expression, regression, disease pattern analysis, pathway enrichment, transcription factor enrichment, and network topology, we identified reproducible gene and pathway signatures associated with increasing NAS and fibrosis severity. Our framework extends previous transcriptomic meta analyses by organizing progression associated signals into biologically coherent modules defined by overlapping pathways and by examining how these modules emerge, persist, and reorganize across increasing histological severity. Integration of meta analysis derived gene scores with pathway gene and protein interaction network topology further enabled the prioritization of central genes within individual biological modules, rather than only across a single global disease network. We made these results available through the open source MASLD-META NETWORK application, which allows researchers to interactively explore and prune networks, calculate alternative centrality metrics, query gene level meta analysis results, and contextualize custom gene sets from human, mouse, or zebrafish through ortholog mapping. Together, this framework provides a stage-resolved and module-aware view of MASLD biology, while offering a reusable resource for evaluating findings from independent clinical cohorts and experimental systems.

Our analysis of clustered inter-pathway gene overlap networks, independent of histological scores, revealed that extracellular matrix remodeling, actin cytoskeleton organization, lipid metabolism, immune system activation and inflammation, and coagulation associated tissue repair constitute the primary biological modules of MASLD progression. These modules align closely with existing literature. Extracellular matrix remodeling remains a core concept in MASLD research, driven by fibrogenesis that leads to the excessive deposition, bundling, and cross linking of collagens (Fan et al., 2024). Furthermore, exposure to fatty acids completely disrupts the cellular mechanosensing machinery in hepatocytes, including actin stress fibers (Rudolph & Chin, 2024). The disease is also fundamentally characterized by hepatic lipid accumulation (Carli et al., 2024), dynamic shifts in liver macrophage populations alongside the accumulation of inflammatory T cells, and hepatocellular damage that compromises the liver’s ability to produce essential coagulation components (He et al., 2025; Pezzino et al., 2024).

Recent transcriptomic studies have demonstrated coordinated changes in immune, fibrotic, and metabolic programmes across MASLD progression. Our meta analytic approach complements these findings by identifying histologically defined comparisons at which coordinated biological modules become reproducibly detectable across independent cohorts (Kamzolas et al., 2026; Pantano et al., 2021). In the NAS analysis, immune and inflammatory dysregulation was evident in the earliest contrast tested, i.e., NAS 3 versus NAS 0 to 2, with modules related to immune and inflammatory dysregulation with oxidative stress and adaptive immune activation and ERK/MAPK signaling. This is consistent with sustained immune pathway activity across the molecular MASLD trajectory and with experimental evidence implicating adaptive immune responses in MASH associated liver injury (Kamzolas et al., 2026; Dudek et al., 2021; Barrow et al., 2021). At NAS 4, however, the dominant network structure shifted toward broader tissue remodelling, including extracellular matrix remodelling, actin cytoskeleton organization, and coagulation associated tissue repair. Along the fibrosis axis, extracellular matrix remodelling and cell adhesion and homeostasis were already detected at F2, whereas adaptive immunity, actin cytoskeleton organization, and coagulation associated tissue repair were detected at F3. This is consistent with transcriptomic evidence showing increasing extracellular matrix associated transcription and expansion of hepatic stellate cell and other nonparenchymal signatures with fibrosis severity, as well as trajectory based evidence placing extracellular matrix remodelling predominantly within mid to late MASLD progression (Pantano et al., 2021; Kamzolas et al., 2026). Metabolic programmes exhibited a more complex organization. Amino acid and nitrogen transfer metabolism was detected at NAS 4, while distinct carbohydrate and insulin response and lipid, bile acid, and steroid metabolic modules became detectable at NAS 5. Along the fibrosis axis, multiple discrete metabolic modules detected at F2 converged into a broader oxidative intermediary metabolism module at F3 and F4. Rather than indicating the biological onset of metabolic dysfunction at these histological scores, these patterns likely reflect changes in the magnitude and coordination of metabolic dysregulation, consistent with the complex, nonmonotonic metabolic dynamics observed across continuous molecular MASLD progression (Kamzolas et al., 2026).

Notably, our NAS comparisons also identified a PI3K/AKT signaling module. This pathway is critical for regulating lipid metabolism, inflammatory responses, and insulin resistance; indeed, previous research demonstrates that impaired PI3K/AKT signaling accelerates MASLD development (Wu et al., 2020). In a related finding, a WNT signaling module emerged in our fibrosis comparisons. Literature indicates that high fat diets induce hepatic steatosis in murine models, and beta catenin stabilization via WNT signaling promotes lipid synthesis through the Akt/mTOR pathway (Wang et al., 2023). Ultimately, comparing modules across histological scoring systems highlights a key biological divergence: fibrosis associated networks captured a higher proportion of development related pathways, whereas NAS associated modules were predominantly driven by inflammation and metabolism.

The availability of harmonized clinical metadata is also important for resolving heterogeneity in MASLD progression. Using datasets with available sex annotations, we extended our framework to sex stratified analyses, which revealed largely conserved module organization but selected differences in coagulation, WNT signaling, and metabolic pathways (Supplementary File 1). More consistent reporting of metadata, including age, will enable broader stratified analyses in future studies.

Through our network analysis, we identified central genes within both biological module networks and STRING derived protein protein interaction networks. Extracellular matrix related genes dominated the most central positions across NAS and fibrosis analyses, including COL3A1, COL1A1, COL1A2, COL5A1, THBS1, THBS2, SPP1, MMP2, and TIMP1. Their convergence across complementary network structures supports extracellular matrix remodelling as a core organizing feature of MASLD progression that connects fibrogenesis, inflammation, and tissue repair. Collagens and MMP2 are established markers of liver fibrosis, while THBS1 promotes fibroblast activation and TGFB1 signalling, and SPP1 links extracellular matrix remodelling with lipid loaded macrophage activity (Seko & Yamaguchi, 2026; Xu et al., 2026; Cho et al., 2026; Dai et al., 2026).

Central genes also included regulators of stromal activation, immune remodelling, and cell fate. TGFB1 and PDGFA represent major fibrogenic growth factors, while PDGFRA and THY1 are associated with activated myofibroblasts (Yang et al., 2025; Colella et al., 2025). The centrality of GPNMB and LGALS3 further supports the contribution of macrophage associated programmes, whereas BCL2, SOX9, and ANXA2 connect disease progression with cell survival, differentiation, and metabolic regulation (Sedda et al., 2025; Wang et al., 2025; Deng et al., 2025; Wu et al., 2024).

MASLD-META NETWORK additionally enabled the systematic reassessment of six previously reported gene panels. Although agreement among the published panels was limited, integration of meta analysis support, network centrality, and biological module coverage consistently prioritized COL1A1, COL3A1, THBS2, FBLN5, and PDGFA across both NAS and fibrosis networks. A stage-resolved expression analysis further identified SULF2, MMP14, IL32, GPNMB, and COL3A1 as altered in the earliest NAS comparison. Notably, these genes encompass complementary processes associated with early liver injury, including inflammatory signalling and insulin resistance through IL32, macrophage associated responses through GPNMB, extracellular matrix remodelling and injury responses through MMP14, and fibrogenic signalling and matrix deposition through SULF2 and COL3A1 (Dali Youcef et al., 2019; Nakamura et al., 2021; Kelly et al., 2024; Wang et al., 2025; Matsumoto et al., 2022). Their detection in the earliest NAS contrast therefore suggests that inflammatory and tissue remodelling programmes may already be transcriptionally engaged at relatively low disease activity, before broader stage associated programmes become detectable at higher NAS values.

Thus, our framework refined heterogeneous published panels into a smaller set of candidates supported by cross study reproducibility, network position, functional coverage, and timing of differential expression.

A particularly important outcome of our work is the identification of LAMA2 and LAMA3 as novel central genes in MASLD progression. While previous research has shown that the laminin gamma 2 chain accelerates lipid accumulation in MASLD (Chen et al., 2025), our findings highlight the alpha subunits as critical central drivers. Given that laminins are known to interact primarily with hepatocytes and liver endothelial cells, these alpha subunits represent promising, novel biomarkers for monitoring MASLD progression (Boel et al., 2025).

As regulators of these central genes and biological modules, we identified SMAD3, RUNX2, RELA, and TP53 as candidate transcriptional regulators of genes associated with MASLD progression. Their enrichment across both NAS and fibrosis analyses suggests that these regulatory programmes are associated with disease severity across complementary histological measures. This finding is consistent with our previous work on TP53 and with experimental evidence linking increased TP53 expression to hepatic fibrosis (Keskus et al., 2026; Kodama et al., 2011). SMAD3 is a central mediator of canonical TGFB1 signalling, which has been implicated in the transition from steatosis to steatohepatitis through hepatocyte injury and subsequently in stellate cell activation and fibrogenesis (Yang et al., 2014; Liu et al., 2019). Together with the high centrality of TGFB1 in our network, enrichment of SMAD3 targets supports this signalling axis as a recurrent regulatory component of MASLD severity. RUNX2 and RELA have likewise been implicated in macrophage associated inflammation and metabolic liver injury (Zhong et al., 2020; He et al., 2025), further supporting the biological relevance of the transcriptional regulators identified across our analyses.

Several limitations to our study must be acknowledged. First, our analyses relied exclusively on bulk transcriptomic data, which precludes the spatial and single cell resolution necessary to map central genes and biological modules to specific hepatic cell subpopulations. Second, the absence of complementary multi omics data restricts our ability to functionally validate these transcriptomic signatures at the proteomic or metabolomic levels, particularly regarding whether our top scoring genes yield detectable, blood based biomarkers for clinical use. Finally, our reliance on cross sectional datasets necessitated defining disease progression through histological scoring. These scores exhibit inherent intra observer and intra stage variability. Moreover, the lack of uniformly reported metadata across the independent cohorts prevented us from systematically controlling for critical clinical confounders, such as patient age, concurrent diabetes, and medication use.

Despite these limitations, our comprehensive meta analysis and the subsequent development of the MASLD-META NETWORK application offer a robust, stage resolved atlas of disease progression. By providing an interactive, open source platform for querying network topology and contextualizing independent experimental data, this framework bridges the gap between computational systems biology and translational hepatology, ultimately accelerating the discovery of novel therapeutic targets and non-invasive diagnostic biomarkers for MASLD.

## Code and Software Availability

All analysis code and data used in this study will be made publicly available upon publication. The MASLD-META NETWORK application can be accessed at https://masld.scilicium.com/.

---

**Supplementary Figure 1.** *Distribution of NAFLD Activity Scores Across Datasets with Proposed Grouping Strategy.* NAS values from zero to eight are grouped into five categories: 0-1-2, 3, 4, 5, 6-7-8. The x-axis lists GEO accession numbers for each individual study, with an additional “Total” column summarizing sample counts across all datasets. The y-axis indicates the number of samples within each NAS group.

**Supplementary Figure 2.** *Sex stratified enrichment of pathways associated with NAS and fibrosis severity*. **A,** Over represented pathways identified separately in female and male participants by comparing low NAS, defined as NAS 0 to 2, with high NAS, defined as NAS 6 to 8. Selected pathways are shown from the coagulation associated tissue repair and metabolic reprogramming modules. **B,** Over represented pathways identified separately in female and male participants by comparing early fibrosis, defined as F0 and F1, with advanced fibrosis, defined as F4. Selected pathways are shown from the WNT signalling and lipid and small molecule metabolism modules. Dot size represents the number of genes contributing to each pathway, and dot colour represents enrichment significance as negative log10 transformed q value. Only pathways with q < 0.05 are shown.

## Supplementary

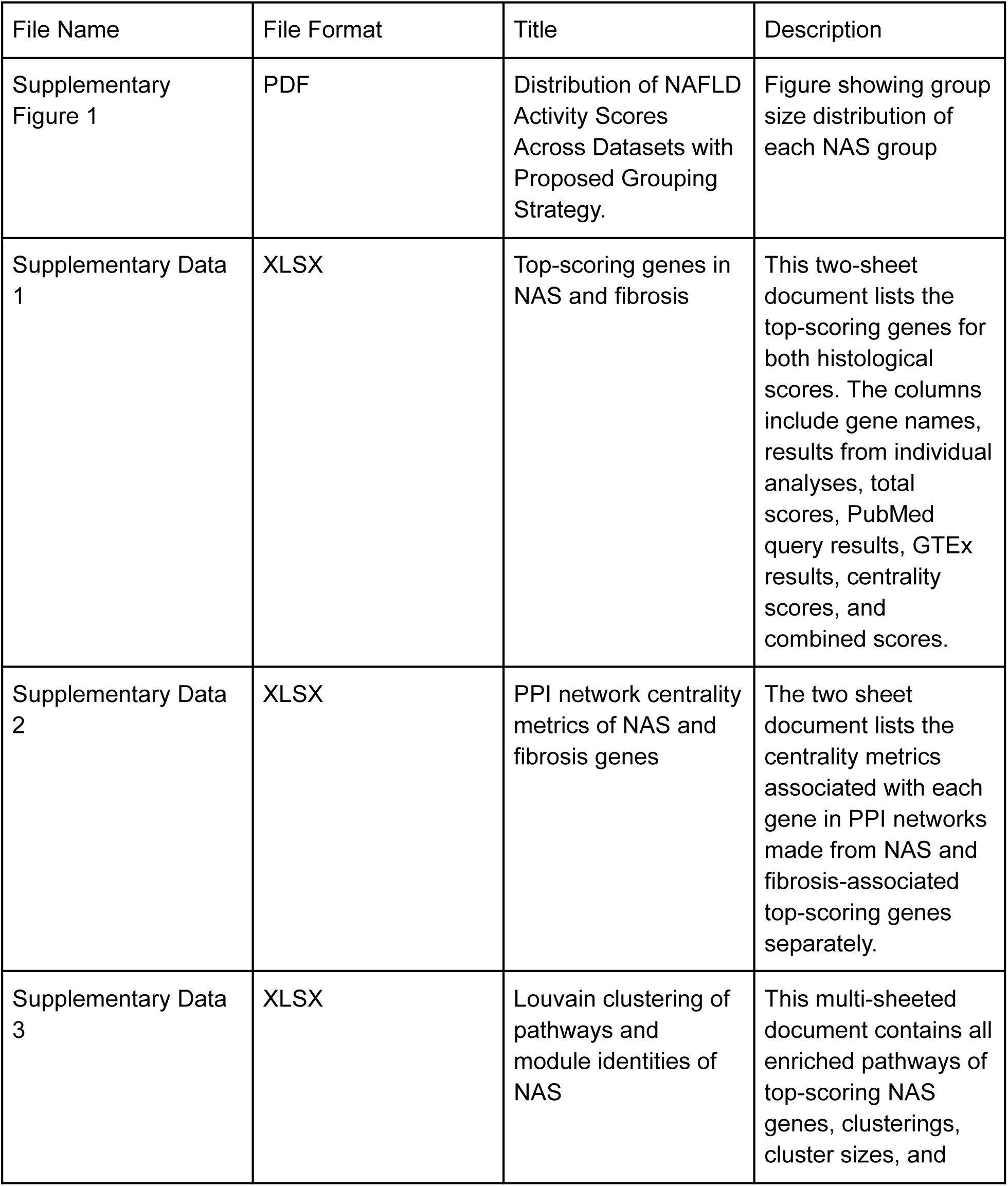
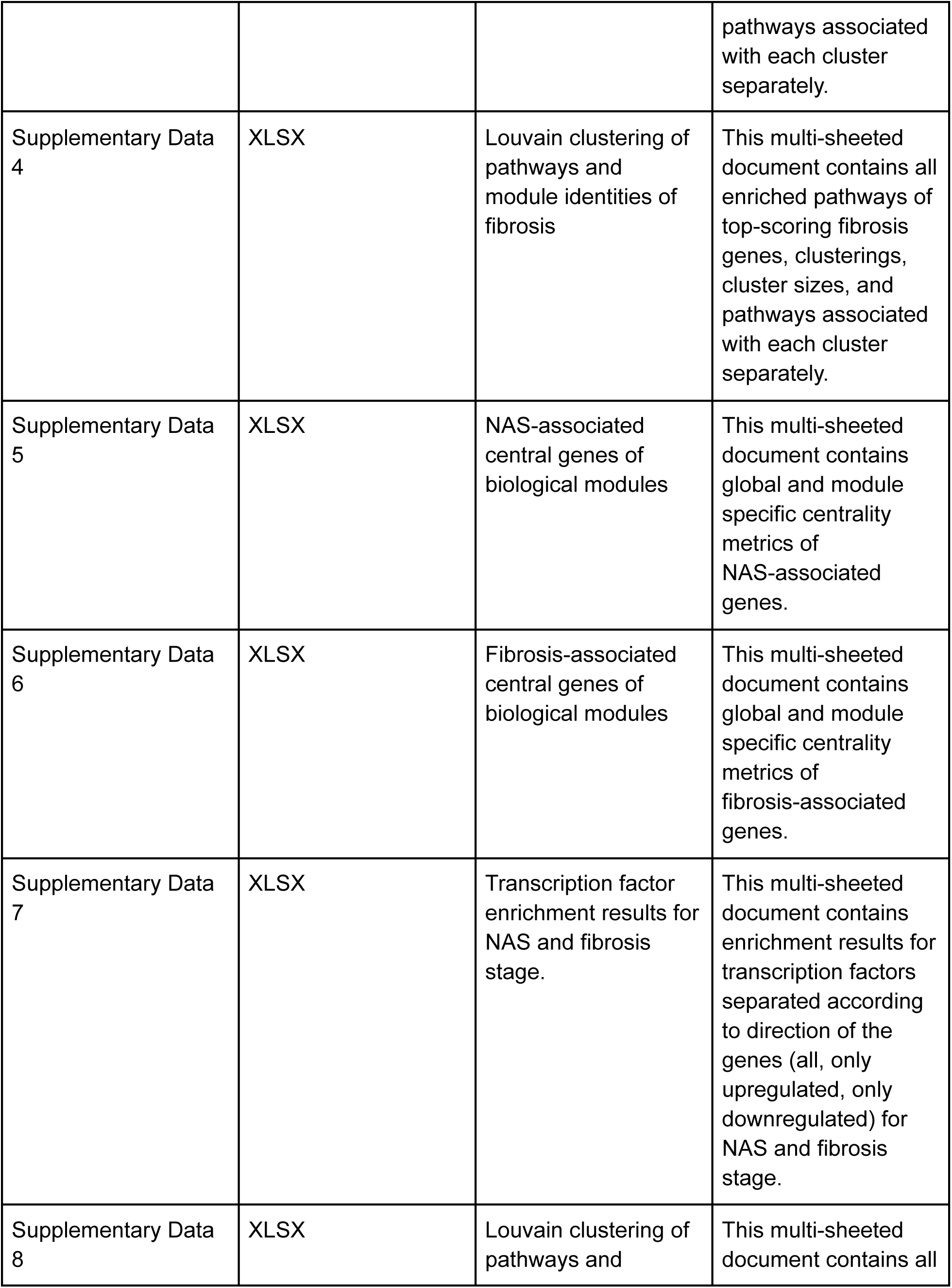
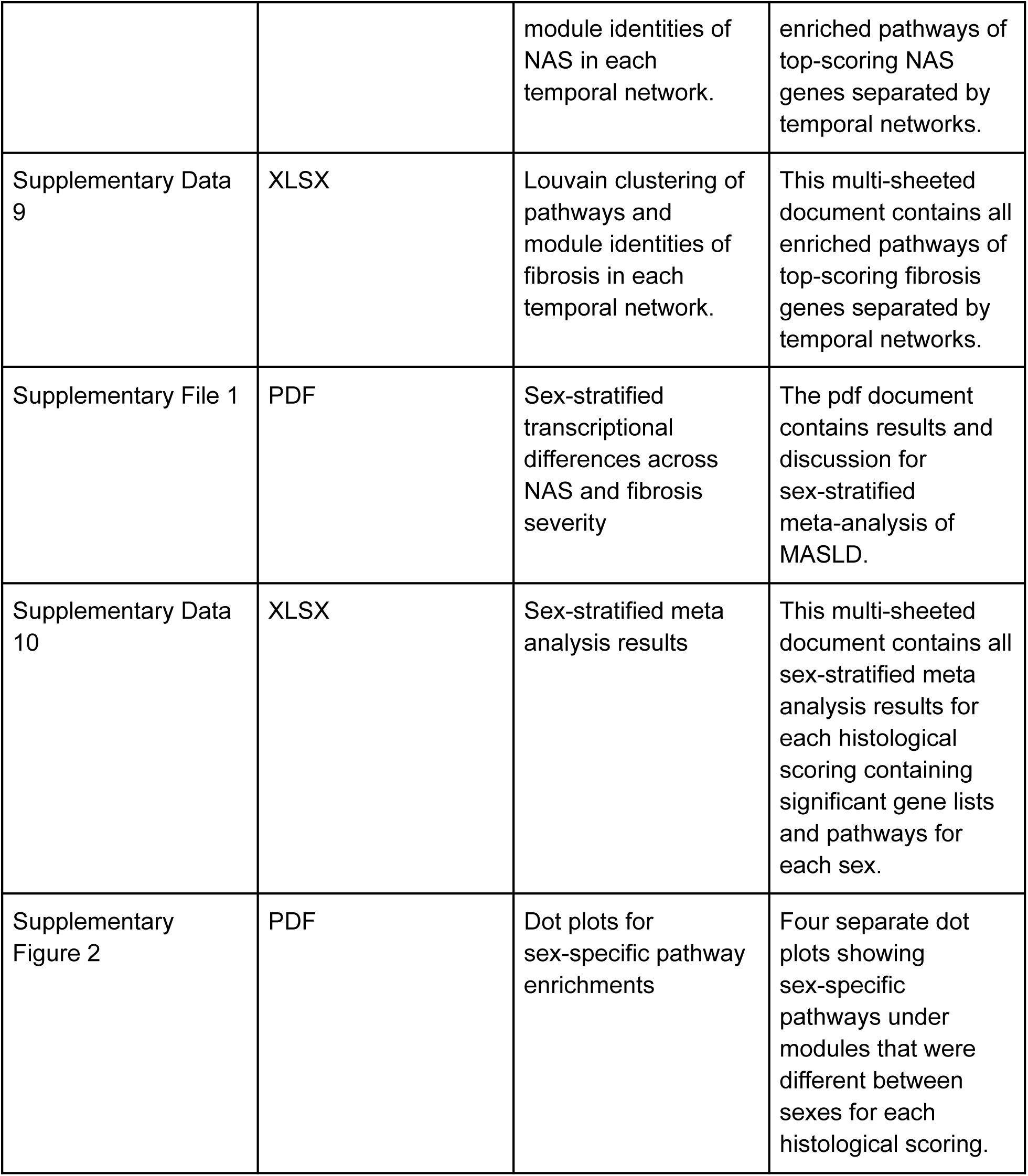

## Notes

### Competing Interest Statement

The authors have declared no competing interest.

### Funding

- European Horizon's research and innovation program HORIZON-HLTH-2022-STAYHLTH-02: 101095679

