## Supplementary figures and images for "Network-based meta-analysis maps stage-dependent molecular programs in MASLD through MASLD-META NETWORK application"

### Supplementary Figure 1

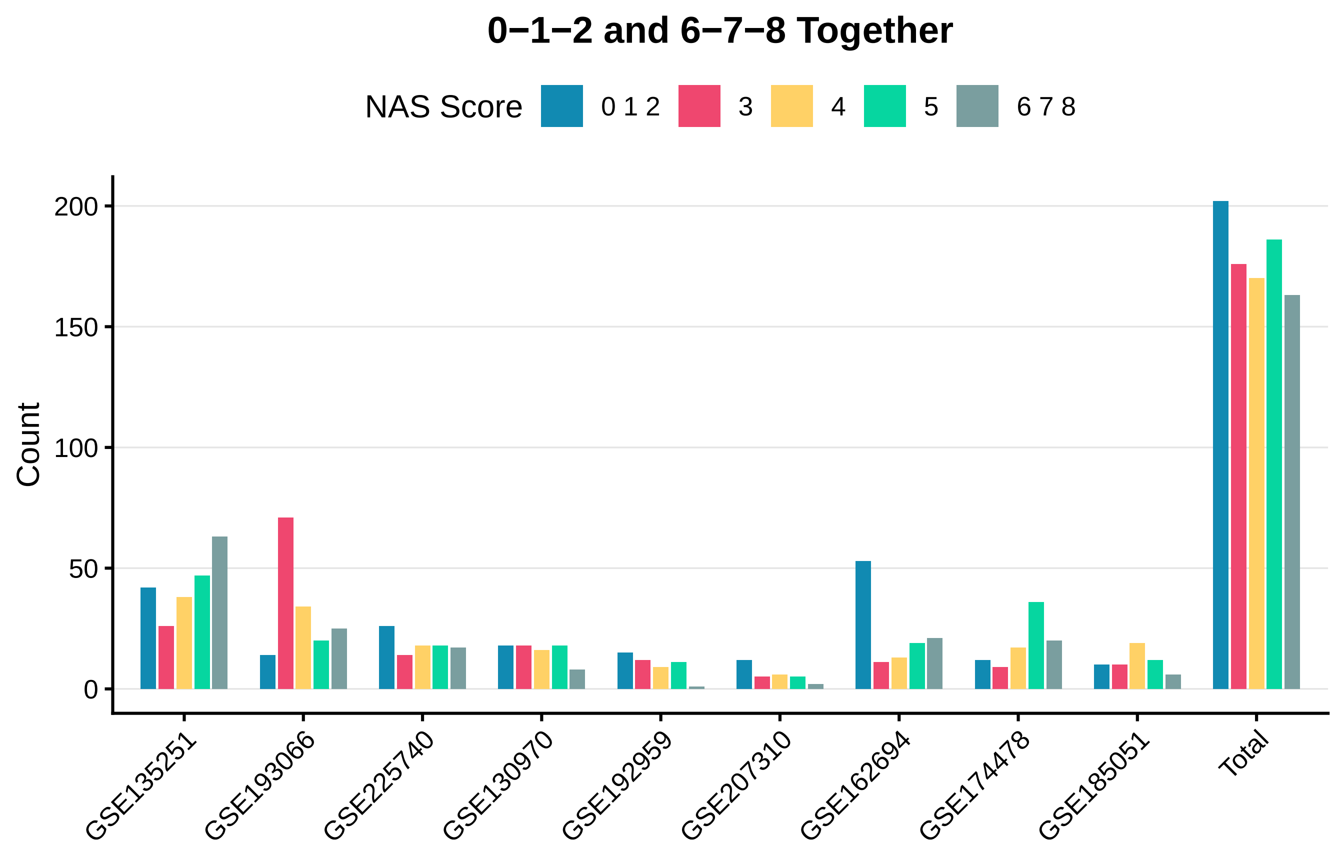

### Supplementary Figure 2

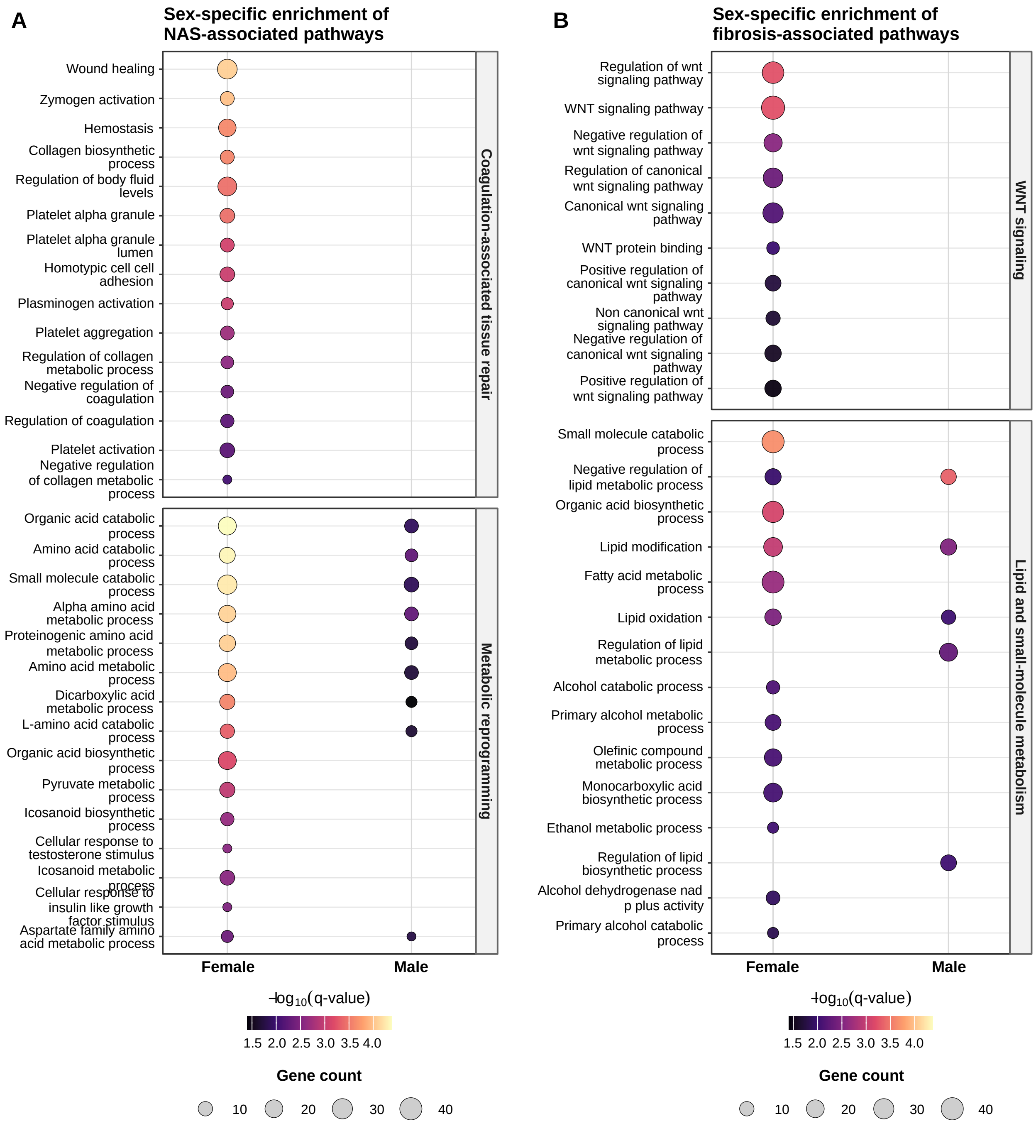
