## Supplementary File 1 for "Network-based meta-analysis maps stage-dependent molecular programs in MASLD through MASLD-META NETWORK application"

### Sex-stratified transcriptional differences across NAS and fibrosis severity

#### RESULTS

To investigate sex-associated differences in MASLD progression, analyses were performed separately in female and male participants by comparing the least severe ($0 \leq NAS \leq 2$) and most severe ($NAS \geq 6$) histological groups in datasets with available sex annotation. Datasets lacking sufficient representation within required comparison groups were excluded from sex-stratified analyses (GSE192959 was excluded from female-based comparisons and GSE207310 from male-based comparisons). Consistent with previous meta-analysis procedures, samples exhibiting irregular p-value distributions were also removed prior to meta-p-value integration.

Following filtering, NAS-based comparisons included GSE130970, GSE207310, GSE162694, GSE174478, and GSE185051 for female participants, and GSE193066 together with GSE174478 for male participants. Genes were retained if they showed $p < 0.05$ in both meta-analysis–derived p-values and were present in the previously defined top-scoring gene lists. Under these criteria, female participants showed a larger number of dysregulated genes (421) compared with male participants (147), with 80 genes shared between sexes in NAS-dependent analyses.

An analogous strategy was applied to fibrosis stage comparisons, contrasting early-stage disease (F0+F1) with advanced fibrosis (F4). After applying identical dataset and gene filtering procedures, female-only fibrosis comparisons included GSE174478, GSE193066, and GSE162694, whereas male-only comparisons included GSE174478, GSE130970, and GSE162694. Similar to the NAS-based analysis, a greater number of dysregulated genes were identified in female participants (995) compared with male participants (523), with 510 genes shared between sexes in fibrosis stage–dependent analyses. The larger difference observed between female and male comparisons in NAS-related analyses relative to fibrosis likely reflects differences in sample size across stratified datasets.

Over-representation analysis was performed across MSigDB gene set collections for sex-stratified gene sets, and C5 ontology gene sets were further analyzed using pathway similarity network approaches as described above. In NAS-dependent analyses, 506 pathways were over-represented in females only, 33 in males only, and 70 were shared between sexes ($q < 0.05$). In the fibrosis stage–dependent analyses, 256 pathways were female-specific, 195 male-specific, and 523 shared between sexes ($q < 0.05$) (Supplementary Data 10).

Comparison of pathway modules identified from NAS-dependent and fibrosis stage–dependent analyses indicated largely conserved modular organization between sexes, with most modules detected in both female and male analyses. Nonetheless, a small number of sex-associated differences were observed. In the fibrosis stage–dependent analyses, a module composed of WNT signaling and regulation of WNT pathway terms (n = 10) was detected only in one sex-stratified analysis (Supplementary Figure 2A). In NAS-dependent analyses, a coagulation-associated tissue repair module comprising 27 pathways showed marked female predominance, with 26 pathways over-represented only in females (Supplementary Figure 2A).

Interestingly, independent of histological scoring system, metabolic process modules in both NAS-dependent and fibrosis stage–dependent analyses included female-specific over-representation of pathways related to fatty acid metabolism, monocarboxylic acid biosynthesis, olefinic compound metabolism, organic acid biosynthesis, and purine nucleotide catabolism.

### DISCUSSION

Another critical factor in MASLD pathogenesis is biological sex. The prevalence of the disease is notably higher in men and postmenopausal women, and previous research has identified significant disparities in the volume of differentially expressed genes and enriched pathways between sexes matched for histological severity (Català-Senent et al., 2021; Kendall et al. 2023) Our analysis revealed that while the majority of our biological modules exhibited no significant sex-specific divergence, females generally presented with a higher number of differentially expressed genes compared to males. The only biological modules demonstrating distinct sex differences were related to coagulation and WNT signaling, which emerged exclusively in our female-specific analyses. Female patients also exhibited more complex metabolic signatures, a finding consistent with previous observations by Kendal et al. However, these results must be interpreted with caution. Sample sizes differed substantially between sexes, particularly in NAS-related analyses. This discrepancy underscores the necessity for future research to consistently include biological sex as metadata in public MASLD datasets. Establishing robust, sex-specific genetic signatures will be paramount for understanding how biological sex impacts disease progression and for refining future diagnostic strategies.
